# Serum nutritional-metabolic recovery during tuberculosis treatment across BMI-defined undernutrition strata: A longitudinal proteomic reanalysis

**DOI:** 10.64898/2026.10.26.761896

**Authors:** Reuben S. Maghembe, Vesla I. Kullaya, Ritah Francis Mutagonda, Megan Null, Peter M. Mbelele, Kennedy M. Ngowi, Jon M. Jacobs, Payam Nahid, W. Evan Johnson, Pranay Sinha, Sana Syed, Stellah George Mpagama, Scott K. Heysell

## Abstract

**Background:** Tuberculosis remains the leading cause of death from a single infectious agent, and undernutrition is a major modifiable risk factor. Serum proteomics can capture systemic host recovery during therapy, but whether baseline BMI-defined undernutrition alters this trajectory is unclear.

**Methods:** We reanalysed processed longitudinal tandem-mass-tag serum proteomics from 30 adults with pulmonary tuberculosis and durable cure sampled at baseline and weeks 8, 17, 26 and 52. Thirteen participants had baseline BMI <18.5 kg/m² and 17 had BMI ≥18.5 kg/m². Protein-level analyses evaluated baseline differences and week-specific within-participant change; module and recovery-axis models evaluated BMI-by-time interactions, with random-intercept mixed models added as a repeated-measures sensitivity analysis. Three complementary treatment-responsive modules were integrated into an exploratory, nonredundant 36-protein nutritional-metabolic summary axis.

**Results:** The reconstructed dataset contained 1,923 quantified features across 150 participant-visits. Inflammatory/acute-phase and metal-handling/nutritional-immunity module scores declined during follow-up, whereas lipid/apolipoprotein-transport, nutritional-carrier and endocrine-immunometabolic scores increased. In the primary week-specific paired-change analyses, no protein-level between-BMI difference in longitudinal change survived false-discovery-rate correction; no curated-module BMI-by-time interaction survived correction. The 36-protein summary axis increased in both BMI strata from week 17 onward. In a random-intercept model without interaction, the BMI <18.5 kg/m² stratum had a lower average axis score across visits than the BMI ≥18.5 kg/m² comparison stratum, although the confidence interval included zero (difference −1.34, 95% CI −2.76 to 0.08; p = .064). Categorical and continuous BMI-by-time interaction tests did not detect differential axis trajectories.

**Conclusions:** Tuberculosis treatment was accompanied by broad longitudinal remodelling of circulating inflammatory and nutritional-metabolic protein systems. An exploratory 36-protein summary axis increased in both baseline BMI strata, while formal interaction tests did not detect BMI-dependent trajectories and remained too imprecise to establish equivalence. BMI should be regarded as an anthropometric proxy rather than a comprehensive measure of nutritional state. The defined axis provides a framework for prospective validation alongside direct measures of body composition, micronutrients, treatment exposure and clinical outcomes.

**Clinical relevance:** Microbiological cure does not necessarily describe the pace or completeness of host recovery from tuberculosis disease. This study summarizes longitudinal lipid-transport, nutritional-carrier and endocrine-immunometabolic changes in a serum recovery axis that can be carried forward for testing in independent cohorts. Baseline BMI was not detectably associated with the axis at baseline in this small cohort and should be regarded as an available anthropometric proxy rather than a comprehensive measure of nutritional state. Future studies should evaluate the axis alongside detailed nutritional phenotyping, pharmacology, post-tuberculosis morbidity and relapse risk.

## 1. Introduction

TB remains a major global cause of morbidity and mortality despite curative antimicrobial therapy. In 2024, an estimated 10.7 million people developed TB and 1.23 million died, making TB again the leading cause of death from a single infectious agent (1, 2). Drug-resistant TB further prolongs treatment and increases toxicity and cost, yet only about two in five people with drug-resistant disease accessed treatment in 2024 (2). Successful therapy must therefore achieve microbial cure while also permitting recovery from host inflammatory, metabolic and tissue consequences of disease.

TB pathobiology reflects sustained interaction between *Mycobacterium tuberculosis* and the host immune-metabolic environment. Macrophage antimicrobial functions span peptide-mediated, metal-handling and phagolysosomal mechanisms (3–5), while membrane and lysosomal injury can reshape host-cell survival and immunometabolism (6, 7). At the tissue level, heterogeneous granuloma microenvironments influence bacterial persistence and treatment response, and inflammatory or structural consequences may persist after bacillary clearance (8, 9).

Undernutrition both increases risk of progression to active TB and can be worsened by disease through anorexia, increased energy expenditure, endocrine disruption and inflammation-driven catabolism (10–12). The World Health Organization estimated that approximately 0.97 million incident TB cases in 2024 were attributable to undernutrition (2). Protein-energy and micronutrient deficits can impair immune and tissue-repair functions, while vitamin and metal biology intersects with acute-phase responses, nutritional immunity and drug disposition (4, 13–15). Conversely, successful treatment can reduce inflammatory consumption and permit restoration of circulating nutritional carriers, lipoproteins and endocrine-metabolic signalling even when body weight changes slowly.

Baseline BMI was treated here as an available anthropometric proxy for undernutrition rather than a comprehensive measure of nutritional state. Although scalable and clinically familiar, BMI does not distinguish lean from fat mass, identify micronutrient deficiencies, resolve hepatic synthetic function or measure immune competence. Pathogen-based assays likewise do not directly report systemic tissue repair or metabolic recovery. Longitudinal serum proteomics can broaden this view, but repeated measures, incomplete observations, batch effects, protein-identifier ambiguity, correlated pathways and extensive multiple testing create analytical challenges (16, 17). Appropriate analysis must therefore preserve time structure, reduce biological redundancy and separate within-person recovery from between-group heterogeneity.

Proteomic and metabolomic studies have shown that active TB and its treatment alter acute-phase, complement/coagulation, extracellular-matrix, apolipoprotein, retinol-carrier and broader metabolic pathways (16, 18–20). These findings support longitudinal serum profiling as a window into host recovery beyond sputum conversion and suggest that lipid transport, nutritional carriers and endocrine signalling are components of the systemic biology linking infection, inflammation and metabolic restoration.

The source dataset, generated by Powell and colleagues, used longitudinal serum from 30 adults with durable cure across five time points to characterize treatment response and recovery beyond the commonly studied 8-week culture-conversion period (17). The original analysis identified extensive time-dependent changes spanning immune, haemostatic, extracellular-matrix and metabolic pathways, together with persistent insulin-like growth factor and lipoprotein-related signals through week 52. However, it was organized primarily around time, treatment regimen and smear-defined severity. It did not test whether baseline BMI-defined undernutrition modified recovery, reconstruct nutrition-relevant protein systems as coordinated modules, or derive a nonredundant integrated nutritional-metabolic recovery measure.

The present reanalysis therefore addressed three linked questions: whether baseline protein abundances or longitudinal changes differed between participants with BMI <18.5 and ≥18.5 kg/m²; whether evidence-guided inflammatory, carrier, lipid, metal-handling, tissue-repair and immune-processing modules showed BMI-dependent trajectories; and whether recurrent lipid-transport, nutritional-carrier and endocrine-immunometabolic signals could be integrated into a robust longitudinal recovery axis associated with baseline BMI or modified by BMI over time.

By combining participant-level reconstruction of the processed TMT dataset, protein-level longitudinal testing, literature-informed module biology, curated identifier remapping, cluster-aware modelling and complementary principal-component constructions, this work presents a BMI-stratified map of serum-proteomic recovery and an exploratory 36-protein nutritional-metabolic summary axis. The axis provides a nonredundant phenotype for testing associations with baseline BMI and BMI-dependent longitudinal modification rather than a direct measure of nutritional status.

## 2. Materials and methods

### 2.1 Study design, source cohort and data governance

This study was a secondary longitudinal reanalysis of de-identified processed serum-proteomics and clinical metadata derived from the MARK-TB data and biospecimen repository and the cohort reported by Powell et al. (17). MARK-TB was established by TB Alliance, the Tuberculosis Trials Consortium (TBTC), and the AIDS Clinical Trials Group (ACTG) to provide biospecimens and associated data in support of tuberculosis biomarker research. The source study included participants enrolled through the REMoxTB trial (21) and the South African National Tuberculosis Program at the same clinical sites. Thirty adults with pulmonary TB and durable cure were randomly selected for the proteomic investigation. The present reanalysis used baseline and weeks 8, 17, 26 and 52, yielding five observations per participant. The original study obtained written informed consent and was approved by the University of California, San Francisco institutional review board (UCSF IRB 12-10360) (17). No new participants, specimens or laboratory experiments were introduced in the present analysis.

The original mass-spectrometry files are publicly deposited in ProteomeXchange/PRIDE under PXD040546 and in MassIVE under MSV000091392. The present work used collaborator-supplied processed abundance, clinical and TMT linkage files that enabled participant-visit reconstruction. These participant-linked processed files were retained in protected local storage and were not redistributed. Public-facing project materials were limited to analysis code, aggregate results, sanitized tables, figures, reference metadata and manuscript text.

### 2.2 Original serum-proteomics generation and processing

The source study collected serum at baseline and weeks 8, 17, 26 and 52. Fourteen high-abundance serum proteins were depleted using a ProteomeLab human IgY14 LC10 affinity column. Proteins were isolated, denatured, digested with trypsin and labelled using 11-plex tandem mass tags (TMT11), with two participants per plex and a pooled reference channel shared across multiplex experiments. Peptides were separated into 12 concatenated offline high-performance liquid-chromatography fractions and analysed on a Thermo Scientific Q Exactive Plus mass spectrometer (17).

Tandem mass spectra were searched with MSGF+ against the April 2017 reviewed human UniProt database. The original workflow controlled false discovery at <0.01% at the unique-peptide level and <0.1% at the protein level. Reporter-ion intensities were summed to protein level, ratioed to the pooled reference within each TMT experiment and log2 transformed (17). The published source analysis reported 1,889 identified proteins. The collaborator-supplied processed matrix used here contained 1,923 unique quantified rows/features and differed numerically from the 1,889 proteins reported in the publication; the available materials did not establish the cause of this difference, and all analyses therefore treated these as quantified matrix features and used unique-protein mappings explicitly where required. After exclusion of 79 sparsely observed features using the ≥10% observed-value criterion, 1,844 protein features were retained in the normalized downstream matrix. No duplicate-protein collapse was assumed.

### 2.3 Participant-visit reconstruction and quality control

TMT channel labels were linked to proteomic matrix columns and then to participant and visit identifiers in the clinical workbook. Reconstruction checks compared participant IDs, scheduled weeks and TMT mappings across files; duplicate participant-visit combinations, unmatched labels and missing scheduled observations were enumerated before analysis. The final matrix contained 150 participant-visit samples from 30 participants, with complete representation at baseline and weeks 8, 17, 26 and 52. The source study’s pooled-reference normalization and log2 transformation were retained. No participant was excluded from the reconstructed longitudinal analysis set, and all were classified as cured.

### 2.4 BMI exposure and analytical questions

Baseline BMI was calculated from the clinical metadata and used as both a categorical and continuous anthropometric proxy. The categorical analysis defined BMI <18.5 kg/m² as undernutrition and BMI ≥18.5 kg/m² as the comparison stratum; this threshold was not treated as evidence that participants above it were nutritionally adequate. The reconstructed cohort included 13 and 17 participants, respectively. The analytical sequence evaluated baseline differences, within-participant change from baseline, between-BMI differences in change and BMI-by-time interaction. Continuous-BMI models were included to avoid relying exclusively on a dichotomous threshold.

### 2.5 Protein-level statistical analysis

Protein-level analyses used log2-normalized abundance values. Baseline group differences were tested with Welch independent-samples t tests. For each follow-up week, a participant-specific delta was calculated as abundance at that week minus baseline abundance. Within each BMI stratum, one-sample t tests assessed whether the mean delta differed from zero. Between-stratum delta differences were evaluated with Welch tests. Benjamini-Hochberg correction controlled the false discovery rate separately within each family of baseline, week-specific between-group and within-group tests (22). Nominal and adjusted p values were retained for complete reporting. These baseline, week-specific between-group change, and within-group change analyses were treated as separate hypothesis families because they addressed distinct inferential questions.

### 2.6 Evidence-guided serum-protein modules and identifier curation

Serum-protein modules were constructed to represent biological programmes relevant to TB pathogenesis, systemic recovery and undernutrition. The framework included inflammatory and acute-phase response; lipid, cholesterol and apolipoprotein transport; retinol and carrier biology; broad nutritional-carrier biology; complement, coagulation and haemostasis; extracellular-matrix and tissue repair; phagocytosis and endocytosis; lysosome and autophagy proxy biology; HLA, antigen-processing and proteasome biology; metal handling and nutritional immunity; vitamin D and humoral immunity; endocrine and systemic immunometabolic signalling; vesicle trafficking; and treatment-response or drug-handling proxy biology. Protein membership was informed by the source proteome, curated gene-protein knowledge and relevant TB, nutrition and immunometabolic literature (4, 6, 13, 14, 16, 17). For module scoring, each protein row was standardized across the 150 samples to a z score, and each module score was the arithmetic mean of available mapped proteins in that sample. Because the processed matrix used UniProt-style mnemonic identifiers for several serum proteins, a curated alias table mapped common mnemonics and alternative labels to gene symbols. The curated analysis reduced duplicate representation and improved coverage of apolipoprotein, retinol-carrier, nutritional-carrier and metal-handling modules. The first-pass and curated analyses were retained as complementary sensitivity layers.

### 2.7 Longitudinal module modelling

Curated module trajectories were analysed with ordinary least-squares models using participant-cluster-robust standard errors. Two model forms were fitted for each module: a continuous-time model testing BMI group by elapsed week, and a categorical-time model testing BMI group by visit week. For the categorical model, baseline was the reference visit and the interaction terms represented BMI-associated differences in change at weeks 8, 17, 26 and 52. Interaction p values were corrected by the Benjamini-Hochberg procedure within the continuous and categorical model families. Module-level descriptive trajectories in Figure 3 show group means with standard errors.

As a post hoc repeated-measures sensitivity analysis, categorical BMI-group-by-week models were additionally fitted using participant random-intercept mixed models. Of the 1,844 retained protein features, 1,834 had sufficient group-by-week support for the full fixed-effect design; the remaining 10 were retained in the audit but were not estimable under this specification. Mixed models were estimated by maximum likelihood (REML = false; L-BFGS optimizer; maximum 500 iterations); when a mixed model failed to converge, ordinary least-squares models with participant-cluster-robust standard errors were used as a computational fallback. Protein interaction p values were adjusted by the Benjamini-Hochberg procedure within each follow-up week, with an additional global correction across all protein-by-week interaction terms. The same categorical sensitivity analysis was applied to the 11 curated modules, with correction across the 44 module-by-week interaction terms.

### 2.8 Exploratory construction and robustness of the serum nutritional-metabolic recovery axis

Three curated modules that showed coordinated treatment-associated behaviour and represented complementary systemic biology were integrated: lipid, cholesterol and apolipoprotein transport; broad nutritional-carrier biology; and endocrine and systemic immunometabolic signalling. Principal component analysis (PCA) was performed on standardized module scores, and the first principal component (PC1) was oriented so that higher values corresponded to increasing values across the selected treatment-responsive modules (23). Because proteins could occur in more than one module, a preferred nonredundant construction was generated from the union of 36 unique proteins. Protein abundances were standardized across samples, remaining missing standardized values were replaced by the feature mean after standardization (approximately zero after centering), the matrix was centred and PC1 was calculated by singular-value decomposition. Axis construction was exploratory rather than prespecified. Because treatment-associated coherence contributed to selection of the three constituent modules, subsequent longitudinal increase of the integrated axis was treated as a descriptive property of the constructed summary measure rather than as an independent test of treatment-associated recovery. The three modules were retained because they combined coordinated treatment-associated behaviour with complementary lipid-transport, carrier, and endocrine/systemic immunometabolic biology; the heterogeneous vitamin C/redox/extracellular-matrix module was retained only as a secondary extension. The 36-protein union contained 134 missing protein-sample values (2.48% of 5,400 cells), confined to APOA5, LEP and LEPR.

Robustness was quantified by Spearman correlations among the three-module score PC1, the 36-unique-protein PC1, the preceding four-module candidate and a unique-protein extension that also contained the heterogeneous vitamin C/redox/extracellular-matrix module. Baseline associations were tested with a Welch test for BMI category and Spearman correlation for continuous BMI. Within-participant change in the preferred axis was assessed at each follow-up week. Standardized paired change (d_z) was calculated as the mean participant-level change divided by the standard deviation of those changes; 95% confidence intervals were generated from 5,000 participant-level bootstrap resamples. Robustness was also evaluated using a 34-protein axis excluding LEP and LEPR and a 33-protein complete-feature axis additionally excluding APOA5; the 33-protein construction required no imputation.

Axis effect modification was evaluated with random-intercept mixed models containing categorical BMI-group-by-week or continuous-BMI-by-week interactions, with participant as the grouping factor (24). Models were estimated by maximum likelihood. When a mixed model did not converge, an ordinary least-squares model with participant-cluster-robust standard errors was used as the model-fitting computational fallback. Interaction p values were adjusted within each model family. A secondary no-interaction random-intercept model estimated the common BMI-group difference across visits while adjusting for categorical visit. Sensitivity models additionally adjusted for baseline sputum smear 3+ or the source-study treatment/disease-category indicator operationally coded as treatcat0.adj = H.

### 2.9 Software and reproducibility

Data reconstruction and statistical analysis were conducted in Python 3.13.2 using pandas for tabular data operations, NumPy for numerical computation and singular-value decomposition, SciPy for statistical tests, and statsmodels for regression, mixed models and multiplicity correction (25–28). Analysis scripts were version-controlled with Git and produced machine-readable tables, figures and run manifests. Private source paths and participant-linked files were excluded from version control. All numerical results in the manuscript were generated from the reconstructed analysis-ready matrix rather than manually transcribed from figures. Generative AI assistance was limited to language editing, structural revision, code/workflow review and submission-formatting support using ChatGPT (OpenAI; version/model GPT-5.6 Sol; source: OpenAI). It was not used to generate proteomic measurements or statistical results; all outputs were reviewed and verified by the authors.

## 3. Results

### 3.1 Reconstructed cohort and longitudinal proteomic matrix

The reconstructed dataset contained 30 participants and 150 participant-visit samples; the collaborator-supplied processed table contained 1,923 unique quantified rows/features, of which 1,844 met the ≥10% observed-value criterion and were retained for normalized downstream analyses. Samples were available at baseline and weeks 8, 17, 26 and 52 for every participant. Thirteen participants had baseline BMI <18.5 kg/m² and 17 had BMI ≥18.5 kg/m² (Table 1). All participants were classified as cured, and no treatment-failure or relapse outcome was available for modelling.

**Table 1.** Baseline characteristics by BMI stratum.

| Characteristic | Overall (N=30) | BMI <18.5 (n=13) | BMI ≥18.5 (n=17) | Absolute SMD |
| --- | --- | --- | --- | --- |
| Age, years | 30.3 (9.4) | 31.7 (8.8) | 29.2 (10.0) | 0.26 |
| Female sex, n/N (%) | 12/30 (40.0%) | 4/13 (30.8%) | 8/17 (47.1%) | 0.34 |
| Source-recorded Black race, n/N (%) | 11/30 (36.7%) | 3/13 (23.1%) | 8/17 (47.1%) | 0.52 |
| Height, cm | 167.2 (11.4) | 166.8 (11.8) | 167.5 (11.4) | 0.06 |
| Weight, kg | 54.7 (11.8) | 46.8 (6.8) | 60.7 (11.5) | 1.47 |
| Baseline BMI, kg/m<sup>2</sup> | 19.4 (3.0) | 16.8 (1.0) | 21.5 (2.3) | 2.63 |
| REMOX study cohort, n/N (%) | 21/30 (70.0%) | 11/13 (84.6%) | 10/17 (58.8%) | 0.60 |
| HIV positive, n/N (%) | 0/30 (0.0%) | 0/13 (0.0%) | 0/17 (0.0%) | 0.00 |
| Cavitation present, n/N (%) | 24/29 (82.8%) | 9/12 (75.0%) | 15/17 (88.2%) | 0.35 |
| Baseline sputum smear 3+, n/N (%) | 11/30 (36.7%) | 8/13 (61.5%) | 3/17 (17.6%) | 1.00 |
| Hard-to-treat disease category, n/N (%) | 9/30 (30.0%) | 6/13 (46.2%) | 3/17 (17.6%) | 0.64 |

Notes: Values are mean (SD) for continuous variables and n/N (%) for categorical variables. Absolute standardized mean differences are descriptive measures of between-stratum imbalance; no baseline hypothesis-test p values are reported. BMI <18.5 kg/m<sup>2</sup> and BMI ≥18.5 kg/m<sup>2</sup> define the exposure strata; therefore baseline BMI and weight were not considered confounders for covariate adjustment. For cavitation, the source sentinel -99 was treated as missing; one participant in the BMI <18.5 stratum lacked cavitation data. Race is reported exactly as the source-recorded Black versus Other/Coloured field and should not be interpreted as a genetically defined category. Treatment arm was not included in the main table because it was structurally unavailable for the nine National TB Program participants. Site was not included in the preferred main table because it was sparse and strongly structured by study cohort; study cohort is retained. All 30 participants were HIV negative, pulmonary-TB cases, baseline culture positive, and ultimately classified as cured in the source analysis set.

**Table 2.**
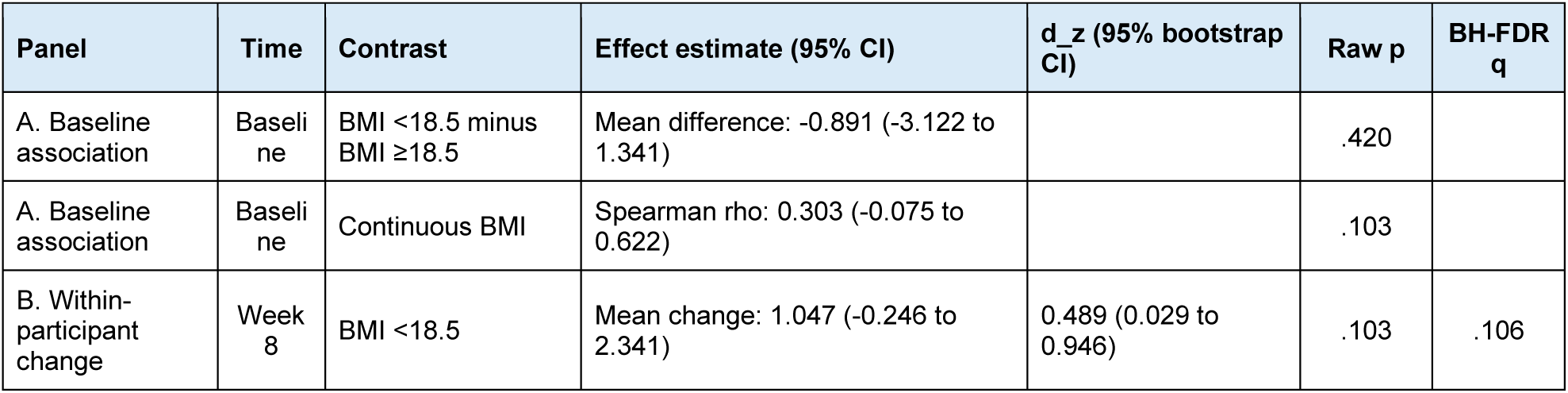
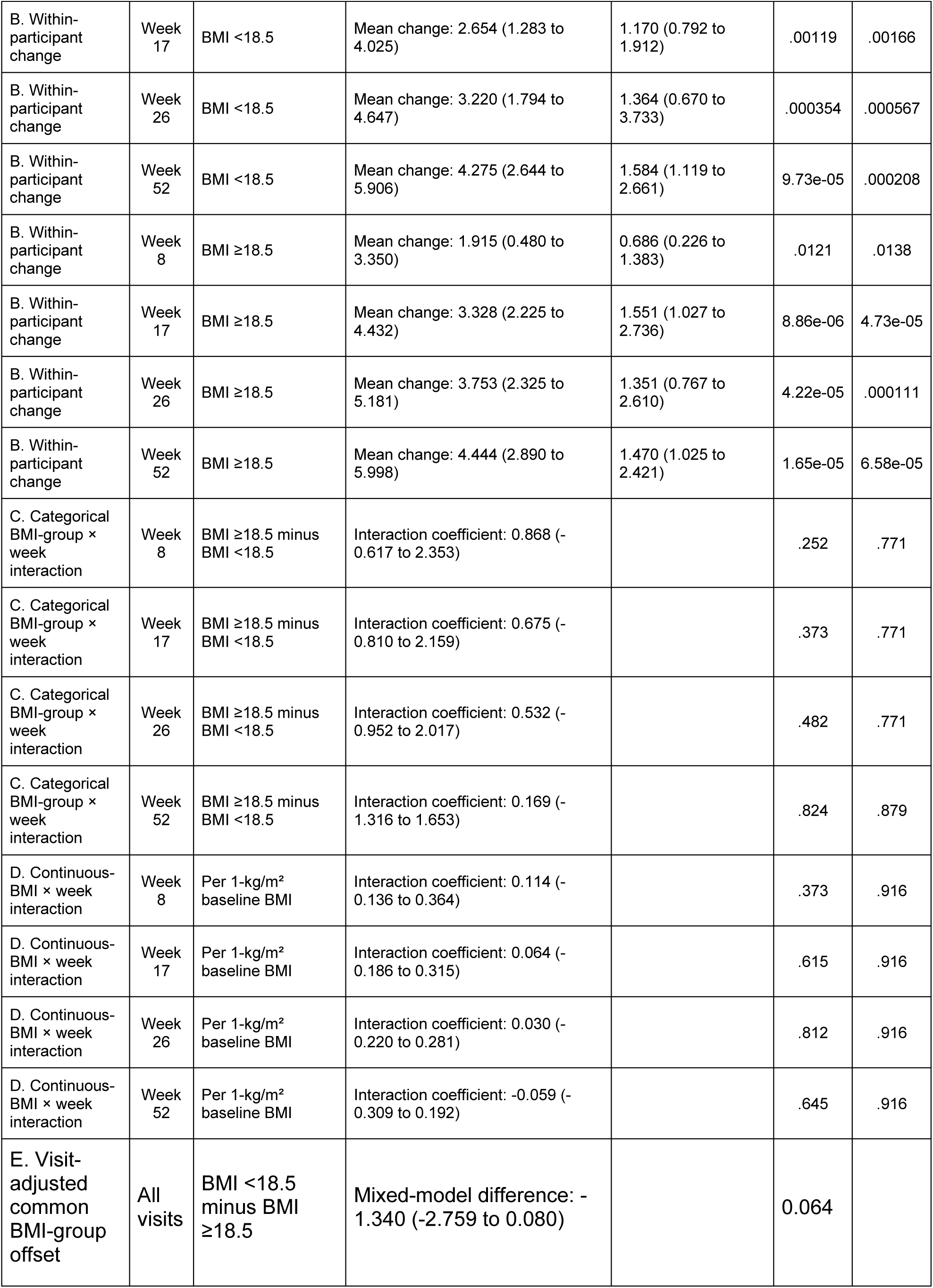
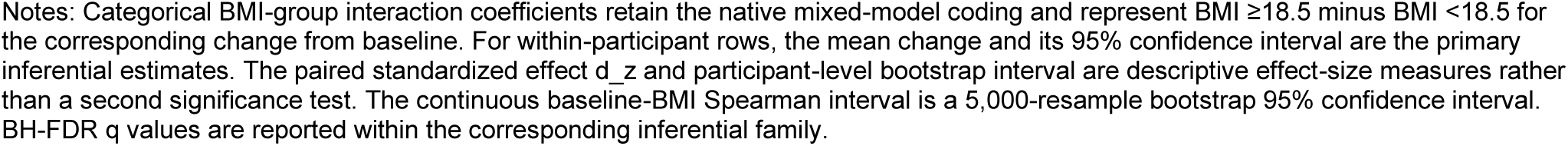
Principal recovery-axis effect estimates and 95% confidence intervals.

| Panel | Time | Contrast | Effect estimate (95% CI) | d_z (95% bootstrap CI) | Raw p | BH-FDR q |
| --- | --- | --- | --- | --- | --- | --- |
| A. Baseline association | Baseline | BMI <18.5 minus BMI ≥18.5 | Mean difference: -0.891 (-3.122 to 1.341) |  | .420 |  |
| A. Baseline association | Baseline | Continuous BMI | Spearman rho: 0.303 (-0.075 to 0.622) |  | .103 |  |
| B. Within-participant change | Week 8 | BMI <18.5 | Mean change: 1.047 (-0.246 to 2.341) | 0.489 (0.029 to 0.946) | .103 | .106 |
| B. Within-participant change | Week 17 | BMI <18.5 | Mean change: 2.654 (1.283 to 4.025) | 1.170 (0.792 to 1.912) | .00119 | .00166 |
| B. Within-participant change | Week 26 | BMI <18.5 | Mean change: 3.220 (1.794 to 4.647) | 1.364 (0.670 to 3.733) | .000354 | .000567 |
| B. Within-participant change | Week 52 | BMI <18.5 | Mean change: 4.275 (2.644 to 5.906) | 1.584 (1.119 to 2.661) | 9.73e-05 | .000208 |
| B. Within-participant change | Week 8 | BMI ≥18.5 | Mean change: 1.915 (0.480 to 3.350) | 0.686 (0.226 to 1.383) | .0121 | .0138 |
| B. Within-participant change | Week 17 | BMI ≥18.5 | Mean change: 3.328 (2.225 to 4.432) | 1.551 (1.027 to 2.736) | 8.86e-06 | 4.73e-05 |
| B. Within-participant change | Week 26 | BMI ≥18.5 | Mean change: 3.753 (2.325 to 5.181) | 1.351 (0.767 to 2.610) | 4.22e-05 | .000111 |
| B. Within-participant change | Week 52 | BMI ≥18.5 | Mean change: 4.444 (2.890 to 5.998) | 1.470 (1.025 to 2.421) | 1.65e-05 | 6.58e-05 |
| C. Categorical BMI-group × week interaction | Week 8 | BMI ≥18.5 minus BMI <18.5 | Interaction coefficient: 0.868 (-0.617 to 2.353) |  | .252 | .771 |
| C. Categorical BMI-group × week interaction | Week 17 | BMI ≥18.5 minus BMI <18.5 | Interaction coefficient: 0.675 (-0.810 to 2.159) |  | .373 | .771 |
| C. Categorical BMI-group × week interaction | Week 26 | BMI ≥18.5 minus BMI <18.5 | Interaction coefficient: 0.532 (-0.952 to 2.017) |  | .482 | .771 |
| C. Categorical BMI-group × week interaction | Week 52 | BMI ≥18.5 minus BMI <18.5 | Interaction coefficient: 0.169 (-1.316 to 1.653) |  | .824 | .879 |
| D. Continuous-BMI × week interaction | Week 8 | Per 1-kg/m<sup>2</sup> baseline BMI | Interaction coefficient: 0.114 (-0.136 to 0.364) |  | .373 | .916 |
| D. Continuous-BMI × week interaction | Week 17 | Per 1-kg/m<sup>2</sup> baseline BMI | Interaction coefficient: 0.064 (-0.186 to 0.315) |  | .615 | .916 |
| D. Continuous-BMI × week interaction | Week 26 | Per 1-kg/m<sup>2</sup> baseline BMI | Interaction coefficient: 0.030 (-0.220 to 0.281) |  | .812 | .916 |
| D. Continuous-BMI × week interaction | Week 52 | Per 1-kg/m<sup>2</sup> baseline BMI | Interaction coefficient: -0.059 (-0.309 to 0.192) |  | .645 | .916 |
| E. Visit-adjusted common BMI-group offset | All visits | BMI <18.5 minus BMI ≥18.5 | Mixed-model difference: -1.340 (-2.759 to 0.080) |  | 0.064 |  |

Notes: Categorical BMI-group interaction coefficients retain the native mixed-model coding and represent BMI $\geq 18.5$ minus BMI $< 18.5$ for the corresponding change from baseline. For within-participant rows, the mean change and its 95% confidence interval are the primary inferential estimates. The paired standardized effect $d_z$ and participant-level bootstrap interval are descriptive effect-size measures rather than a second significance test. The continuous baseline-BMI Spearman interval is a 5,000-resample bootstrap 95% confidence interval. BH-FDR q values are reported within the corresponding inferential family.

**Table 3.** Preferred 36-protein recovery-axis membership and PC1 loadings.

| Protein feature | Source module membership | PC1 loading |
| --- | --- | --- |
| AFAM_HUMAN | Broad nutritional-carrier | 0.2768 |
| ALBU_HUMAN | Endocrine/immunometabolic; Broad nutritional-carrier | 0.0891 |
| APOA1_HUMAN | Endocrine/immunometabolic; Broad nutritional-carrier;<br>Lipid/apolipoprotein transport | 0.2119 |
| APOA2_HUMAN | Endocrine/immunometabolic; Broad nutritional-carrier;<br>Lipid/apolipoprotein transport | 0.1260 |
| APOA4_HUMAN | Broad nutritional-carrier; Lipid/apolipoprotein transport | 0.1632 |
| APOA5_HUMAN | Lipid/apolipoprotein transport | 0.0621 |
| APOB_HUMAN | Lipid/apolipoprotein transport | 0.2032 |
| APOC1_HUMAN | Lipid/apolipoprotein transport | 0.2252 |
| APOC2_HUMAN | Lipid/apolipoprotein transport | 0.1781 |
| APOC3_HUMAN | Endocrine/immunometabolic; Lipid/apolipoprotein transport | 0.1336 |
| APOC4_HUMAN | Lipid/apolipoprotein transport | 0.2002 |
| APOD_HUMAN | Lipid/apolipoprotein transport | 0.1768 |
| APOE_HUMAN | Lipid/apolipoprotein transport | 0.1198 |
| APOF_HUMAN | Lipid/apolipoprotein transport | 0.0955 |
| APOH_HUMAN | Lipid/apolipoprotein transport | 0.2640 |
| APOL1_HUMAN | Lipid/apolipoprotein transport | 0.1104 |
| APOM_HUMAN | Lipid/apolipoprotein transport | 0.1961 |
| CETP_HUMAN | Lipid/apolipoprotein transport | 0.1652 |
| CLUS_HUMAN | Lipid/apolipoprotein transport | 0.2296 |
| IBP3_HUMAN | Endocrine/immunometabolic | 0.2713 |
| IBP4_HUMAN | Endocrine/immunometabolic | 0.1340 |
| IBP5_HUMAN | Endocrine/immunometabolic | 0.1225 |
| IBP6_HUMAN | Endocrine/immunometabolic | 0.1373 |
| IGF1_HUMAN | Endocrine/immunometabolic | 0.1592 |
| IGF2_HUMAN | Endocrine/immunometabolic | 0.1457 |
| LCAT_HUMAN | Lipid/apolipoprotein transport | 0.0003 |
| LEPR_HUMAN | Endocrine/immunometabolic | -0.0270 |
| LEP_HUMAN | Endocrine/immunometabolic | 0.1126 |
| PLTP_HUMAN | Lipid/apolipoprotein transport | 0.0849 |
| PON1_HUMAN | Lipid/apolipoprotein transport | 0.1546 |
| PON3_HUMAN | Lipid/apolipoprotein transport | 0.1821 |
| RET4_HUMAN | Endocrine/immunometabolic; Broad nutritional-carrier | 0.2635 |
| SHBG_HUMAN | Endocrine/immunometabolic | 0.1100 |
| TRFE_HUMAN | Broad nutritional-carrier | -0.0349 |
| TTHY_HUMAN | Endocrine/immunometabolic; Broad nutritional-carrier | 0.1242 |
| VTDB_HUMAN | Broad nutritional-carrier | 0.1990 |

### 3.2 Protein-level baseline and longitudinal tests

At baseline, 89 protein rows had nominal BMI-group p values < .05, but none remained significant after Benjamini-Hochberg correction. Week-specific comparisons of participant-level change from baseline identified no protein with a significant between-BMI difference after correction. Within each BMI stratum, numerous proteins changed significantly from baseline during treatment and follow-up. Representative strong responders included decreases in CRP and AACT in both strata, an increase in APOA4 in the BMI ≥18.5 kg/m² stratum, and increases in LUM, COMP, ICOSL and MMP2 in the BMI <18.5 kg/m² stratum.

Figure 2 summarizes the top-ranked within-stratum responders; full ranked results are provided in Supplementary Table S20.

**Figure 1.**
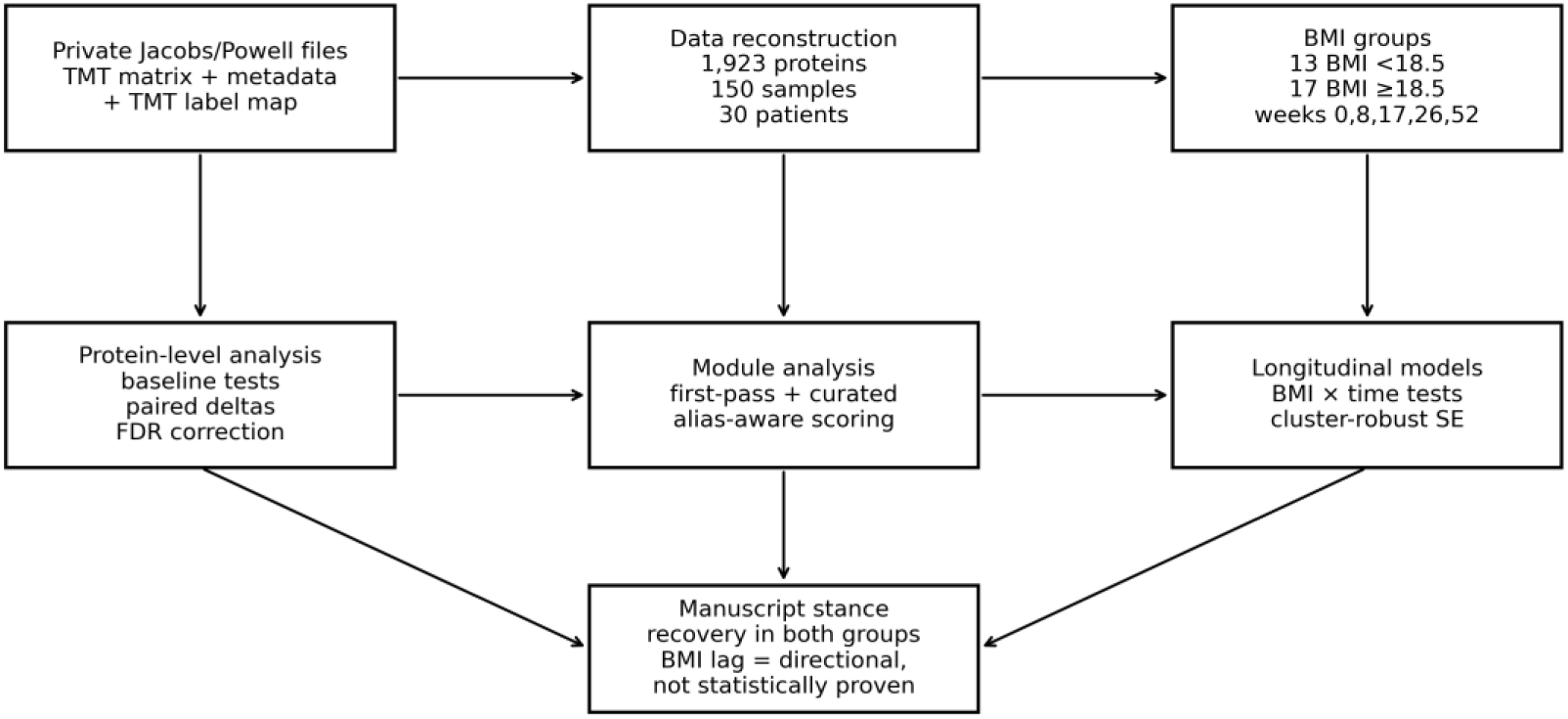
The workflow links collaborator-supplied processed TMT serum-proteomics, clinical metadata and channel mapping to participant-level reconstruction, protein testing, evidence-guided module scoring, BMI-stratified longitudinal models and manuscript-level synthesis.

**Figure 2.**
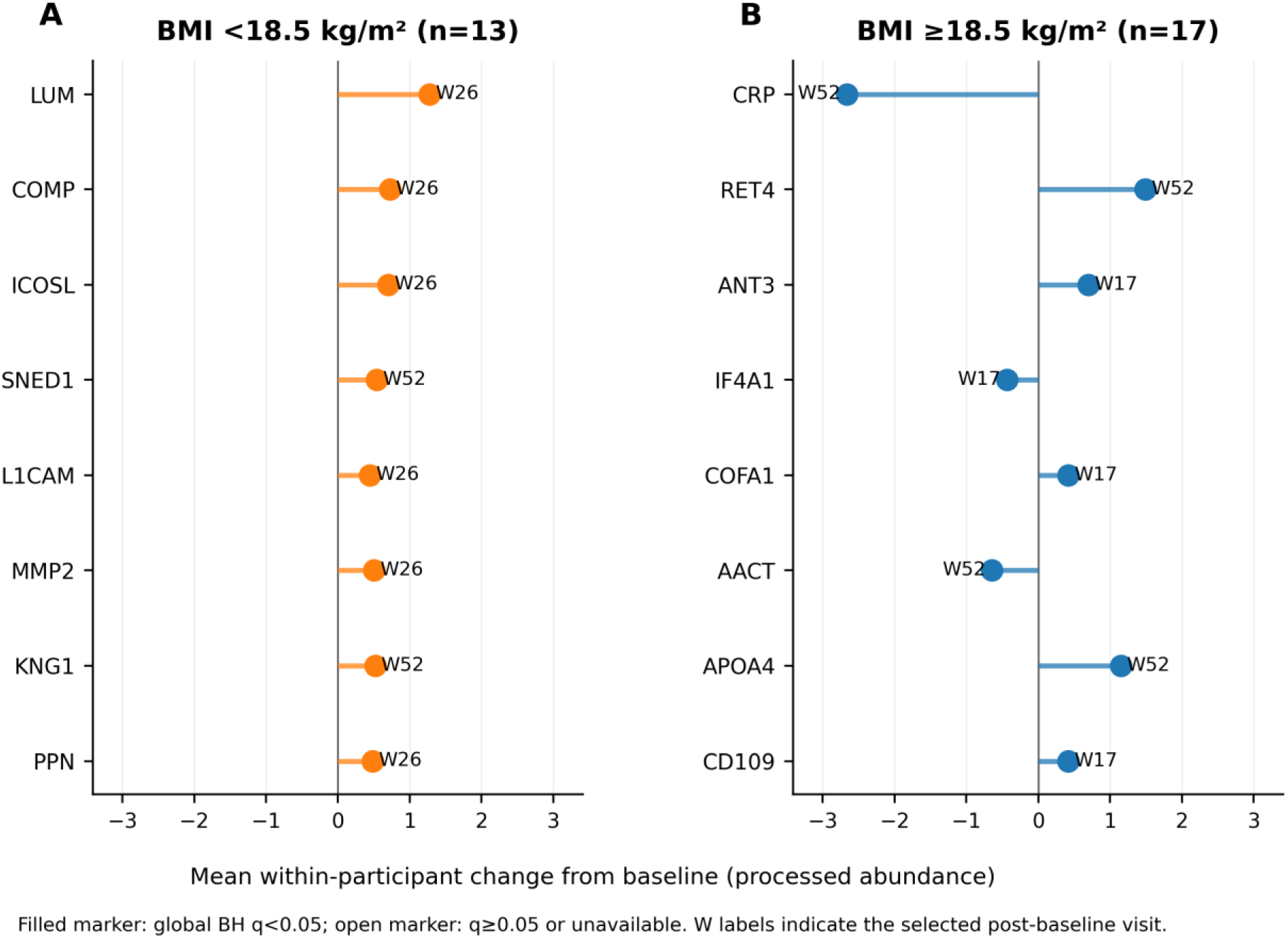
Top-ranked within-stratum protein responses during tuberculosis treatment and follow-up. The eight highest-ranked unique protein responders are shown separately for participants with baseline BMI <18.5 kg/m² (A) and BMI ≥18.5 kg/m² (B). Ranking used the smallest existing global Benjamini-Hochberg-adjusted q value across post-baseline visits for each protein within each stratum, with ties resolved by larger absolute mean change. Points show mean within-participant change from week 0 at the selected visit; W labels identify that visit. Filled markers denote global BH q < .05 and open markers denote q ≥ .05 or unavailable. Selection is descriptive and introduces no additional hypothesis test. Complete ranked results are provided in Supplementary Table S20.

**Figure 3.**
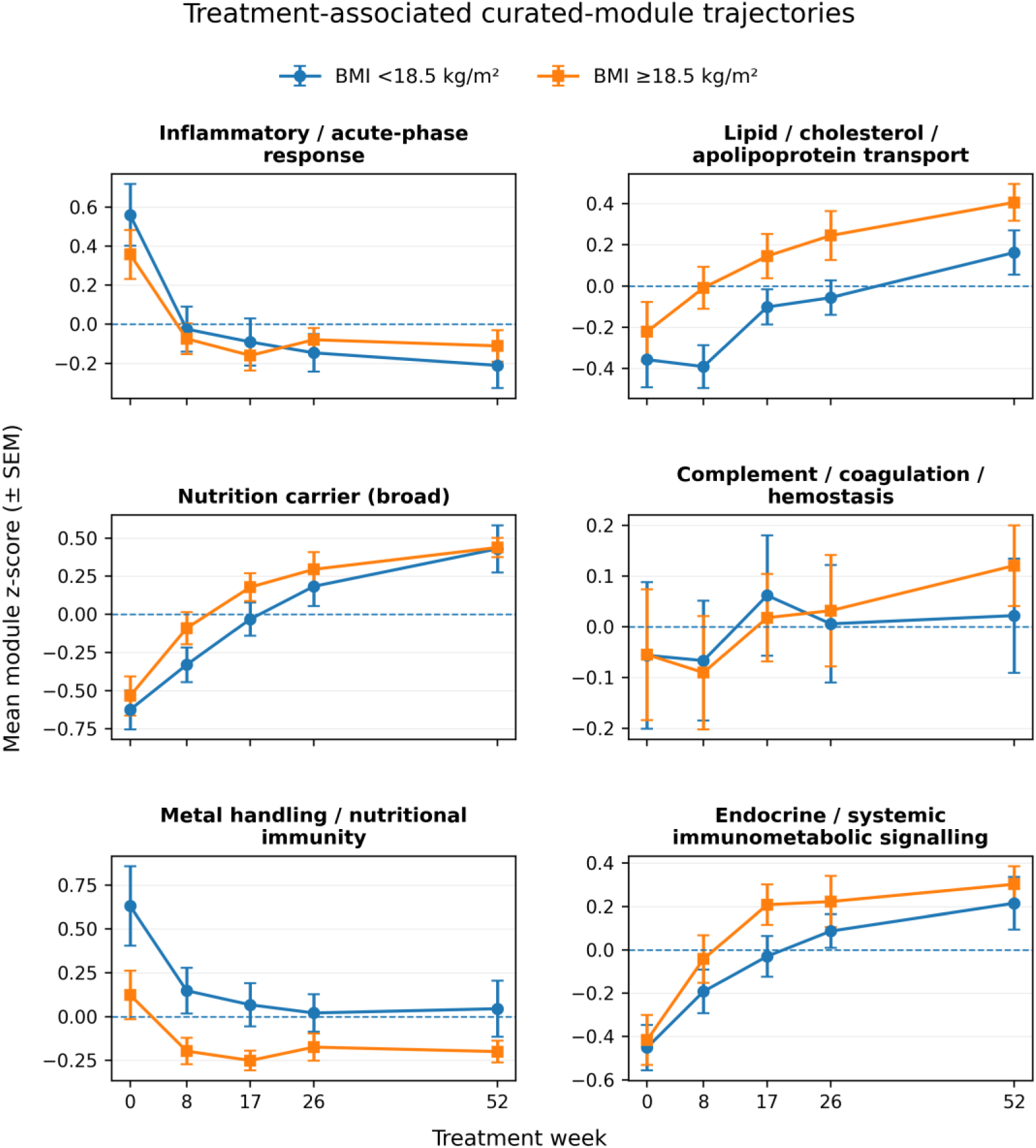
Curated-module trajectories during tuberculosis treatment and follow-up. Mean standardized module scores are shown at weeks 0, 8, 17, 26 and 52 by baseline BMI stratum; error bars indicate standard errors of the mean. The six panels display inflammatory/acute-phase, lipid/apolipoprotein-transport, broad nutrition-carrier, complement/coagulation, metal/nutritional-immunity, and endocrine/systemic immunometabolic modules. The figure is descriptive; formal BMI-group-by-time interaction estimates are presented in Figure 5.

In the post hoc repeated-measures sensitivity analysis, 1,753 of 1,834 protein models converged as participant random-intercept mixed models and 81 used the prespecified participant-cluster-robust OLS fallback. Forty-nine week-specific interaction terms met the within-week Benjamini-Hochberg threshold, and 48 remained significant in the additional global BH audit. However, 41 of the 49 terms were supported by fewer than 10 paired baseline-follow-up participants, none of the corresponding primary paired-change contrasts survived false-discovery-rate correction, and the median mixed-model-to-Welch standard-error ratio among these 49 terms was 0.733. These signals were therefore treated as model-sensitive secondary findings rather than primary protein-level discoveries (Supplementary Tables S21-S23 and S25).

### 3.3 Module-level longitudinal patterns and between-BMI comparisons

The first-pass framework mapped 168 protein rows to serum-proteomic modules. Across follow-up, inflammatory/acute-phase and metal-handling/nutritional-immunity module scores decreased overall, whereas lipid/apolipoprotein-transport, nutritional-carrier and endocrine/systemic-immunometabolic scores increased; complementary retinol-carrier signals also increased. These directional changes were observed in both baseline BMI strata, although not every stratum-specific module-week comparison was false-discovery-rate significant (Figure 3). No between-BMI module-delta comparison was nominally significant or significant after false-discovery-rate correction at weeks 8, 17, 26 or 52 (Figure 4).

**Figure 4.**
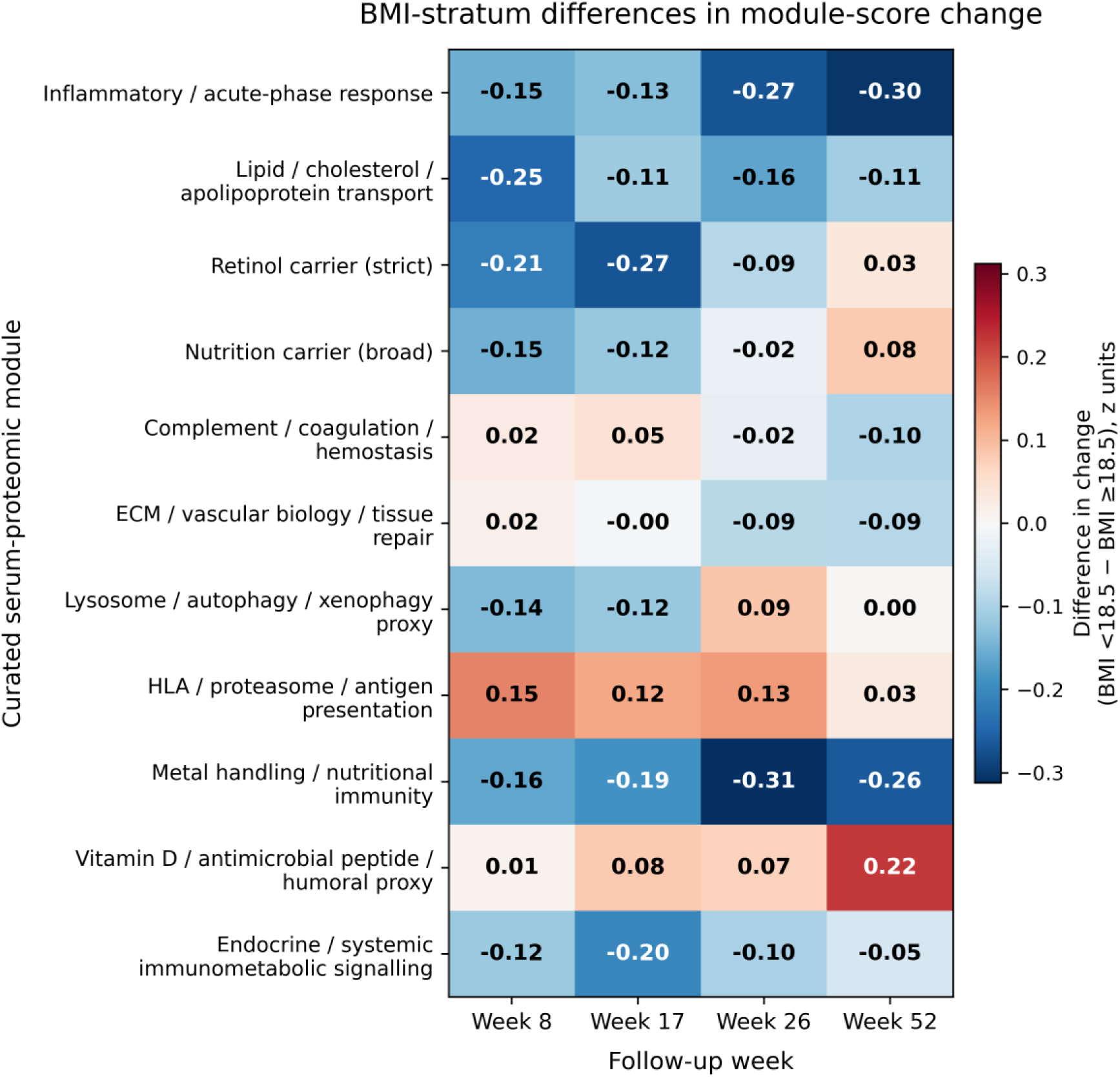
Curated-module delta heatmap comparing baseline BMI strata. Cells show the between-stratum difference in change from baseline, calculated as BMI <18.5 minus BMI ≥18.5 kg/m², for each curated module and follow-up week. The diverging color scale is centered at zero, and cell values report standardized module-score differences. No between-BMI module-delta comparison reached nominal significance or significance after Benjamini-Hochberg false-discovery-rate correction; the directional patterns are therefore hypothesis-generating.

Curated alias-aware remapping produced 11 analysis modules containing 110 unique mapped proteins. The curated trajectories retained time-dependent changes in both BMI strata. Between-BMI delta tests remained non-significant for every curated module and follow-up week (Supplementary Tables S6-S9).

### 3.4 BMI-by-time interaction tests

Continuous-time models fitted 11 BMI-group-by-time interaction tests, and categorical-time models fitted 44 week-specific interaction tests. None of the continuous or categorical interactions had a nominal p value < .05, and none was significant after Benjamini-Hochberg correction (Figure 5; Supplementary Table S10). The smallest interaction p values occurred in vitamin D/humoral-proxy, inflammatory acute-phase, metal/nutritional-immunity, endocrine-immunometabolic and lipid/apolipoprotein modules.

**Figure 5.**
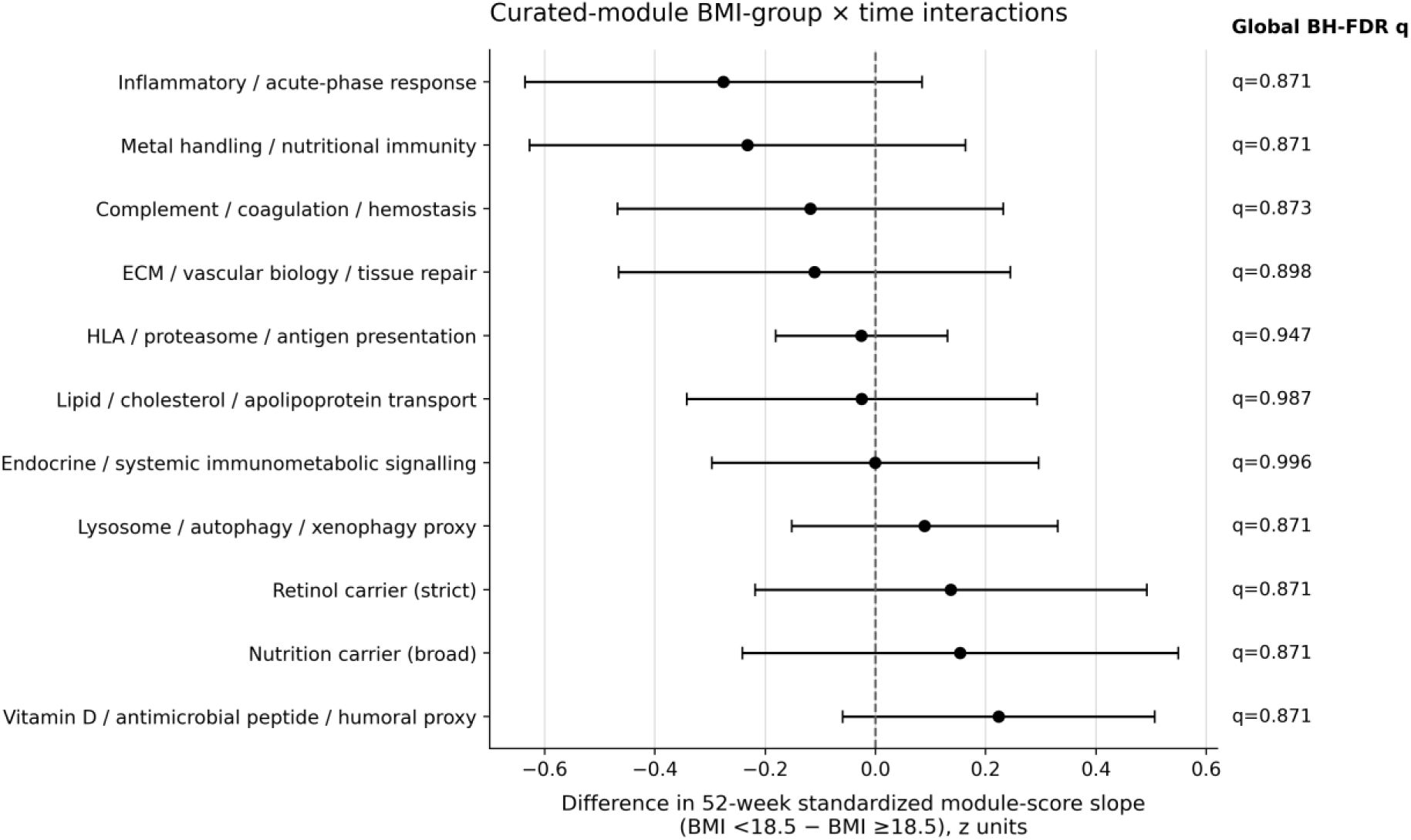
Curated-module BMI-group by continuous-time interaction estimates. Points and horizontal lines show cluster-robust OLS interaction coefficients and 95% confidence intervals for continuous time (visit week/52) by baseline BMI group, with BMI ≥18.5 kg/m² as the reference group and BMI <18.5 kg/m² encoded as 1. Coefficients represent the difference in standardized module-score slope over 52 weeks for BMI <18.5 versus BMI ≥18.5 kg/m². Positive estimates indicate a more positive or less negative slope in the lower-BMI group; negative estimates indicate a less positive or more negative slope. Benjamini-Hochberg false-discovery-rate correction was applied globally across all 55 curated-module interaction tests (11 continuous-time and 44 categorical-week tests); global q values are shown at right. All 11 continuous-time 95% confidence intervals included zero, and none of the 55 tests was nominally significant or significant after FDR correction. The 44 categorical-week interaction estimates are retained in Supplementary Table S10.

In the corresponding categorical repeated-measures sensitivity analysis of the 11 curated modules, eight random-intercept mixed models converged and three used the participant-cluster-robust OLS fallback. None of the 44 BMI-group-by-week interaction terms survived Benjamini-Hochberg correction (Supplementary Table S24).

### 3.5 Recovery-axis construction and robustness

PCA of the three curated module scores produced PC1 loadings of .595 for endocrine and systemic immunometabolic signalling, .571 for broad nutritional-carrier biology and .566 for lipid, cholesterol and apolipoprotein transport. The component explained 86.5% of variance among the three module scores (Figure 6A-B).

**Figure 6.**
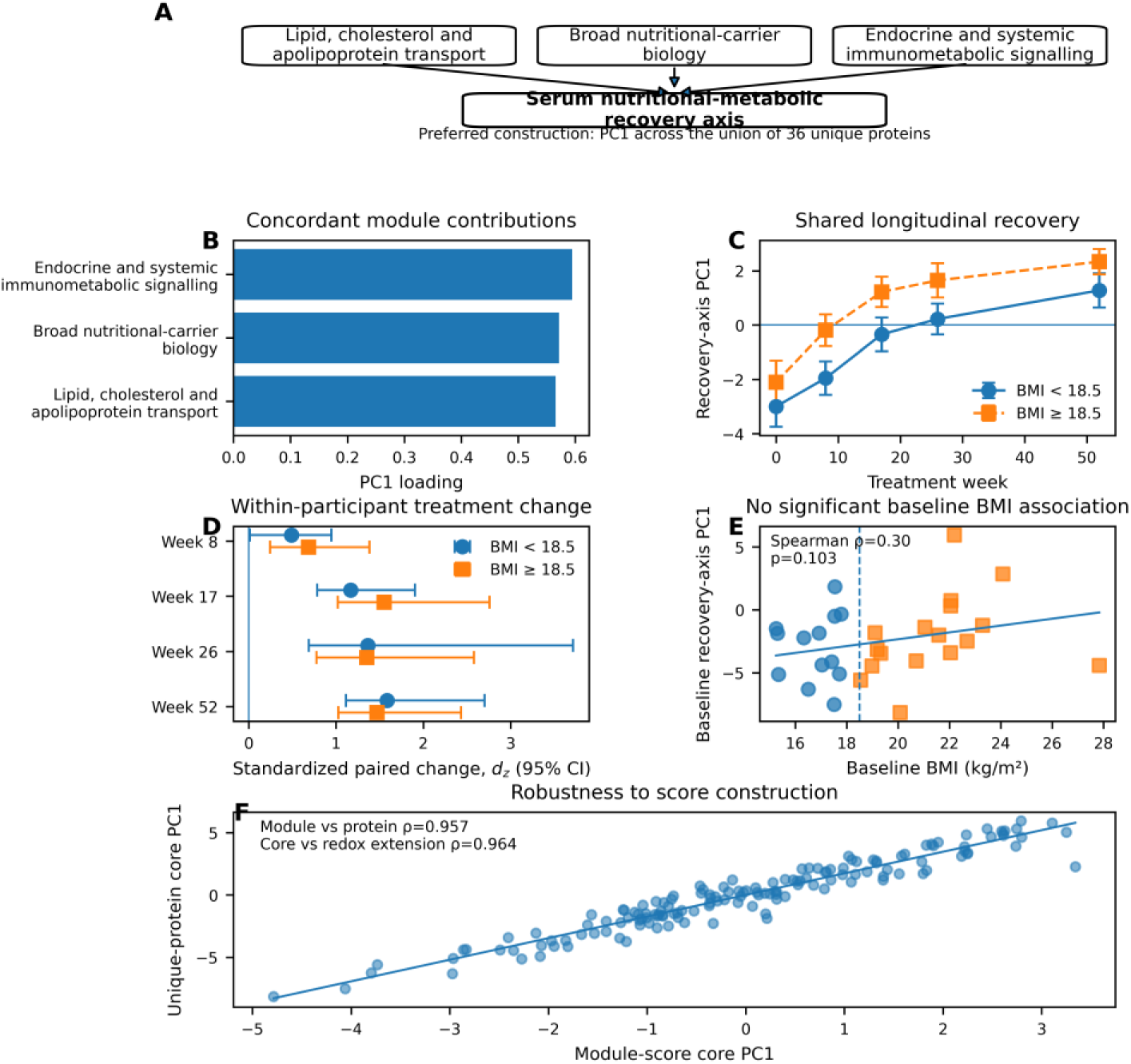
A, Three-module architecture of the integrated axis. B, Module-level PC1 loadings. C, Mean preferred 36-protein recovery-axis score by treatment week and baseline BMI group, with standard errors. D, Standardized within-participant change from baseline with bootstrap 95% confidence intervals. E, Baseline BMI versus baseline recovery-axis score. F, Agreement between module-score and unique-protein PC1 constructions. The axis increased during treatment in both BMI strata; baseline BMI associations and BMI-by-time interactions were not significant.

The preferred nonredundant construction contained 36 unique proteins and its PC1 explained 24.0% of protein-level variance. It correlated strongly with the three-module score PC1 (Spearman ρ = .957), while alternative sensitivity constructions yielded concordant conclusions (Figure 6F; Supplementary Tables S13-S14 and S17).

### 3.6 Longitudinal recovery-axis changes and baseline BMI associations

Among participants with BMI <18.5 kg/m², standardized within-participant change in the preferred axis was d_z = 0.489 at week 8 (BH-adjusted p = .106), 1.170 at week 17 (p = .00166), 1.364 at week 26 (p = .000567) and 1.584 at week 52 (p = .000208). Among participants with BMI ≥18.5 kg/m², the corresponding values were 0.686 at week 8 (p = .0138), 1.551 at week 17 (p = 4.73 × 10−5), 1.351 at week 26 (p = .000111) and 1.470 at week 52 (p = 6.58 × 10−5) (Figure 6C-D; Supplementary Table S15).

Categorical BMI-group-by-week interactions were not significant after correction (all BH-adjusted p ≥ .771), and continuous-BMI-by-week interactions were not significant (all BH-adjusted p = .916) (Supplementary Table S16). The categorical interaction estimates were nevertheless imprecise: the week-8 coefficient was 0.868 (95% CI −0.617 to 2.353) and the week-52 coefficient was 0.169 (95% CI −1.316 to 1.653). Thus, absence of statistically significant interaction should not be interpreted as evidence of equivalent trajectories. At baseline, the mean axis score was −2.998 in the BMI <18.5 kg/m² stratum and −2.107 in the BMI ≥18.5 kg/m² comparison stratum (Welch p = .420); the baseline correlation between continuous BMI and the axis was ρ = .303 (p = .103) (Figure 6E; Supplementary Table S17).

In a random-intercept model without a BMI-by-time interaction, the BMI <18.5 kg/m² stratum had a lower visit-adjusted average recovery-axis score than the BMI ≥18.5 kg/m² comparison stratum (difference - 1.340, 95% CI -2.759 to 0.080; p = 0.064). Sensitivity adjustment for baseline sputum smear 3+ did not attenuate this offset (difference -1.866, 95% CI -3.401 to -0.331; p = 0.017), nor did adjustment for the source-study treatment/disease-category indicator (difference -1.648, 95% CI -3.097 to -0.199; p = 0.026). Existing interaction-sensitivity models likewise showed no detectable BMI-by-time interaction after adjustment for baseline sputum smear 3+ or baseline disease-severity category (Supplementary Table S18). These observational sensitivities do not establish an independent nutritional effect or exclude residual confounding.

## 4. Discussion

Across one year of observation, participants who achieved durable cure showed extensive serum-proteomic change, including increasing scores on an exploratory lipid-transport, nutritional-carrier and endocrine-immunometabolic summary axis. Because treatment-associated behaviour contributed to selection of its constituent modules, the temporal increase itself should not be interpreted as independent validation of a recovery programme. The more informative questions are whether the summary phenotype differs between clinically defined groups and whether its longitudinal trajectory is modified by baseline characteristics.

The broad within-participant proteomic shifts can be understood biologically as the circulation moving away from the acute infectious state and toward renewed homeostatic transport, synthetic and regulatory functions. Active TB drives hepatic acute-phase production, complement and coagulation activation, leukocyte recruitment, endothelial perturbation and extracellular-matrix turnover. These processes redistribute amino acids, lipids, retinoids and metal-binding proteins and can suppress negative acute-phase carriers. As bacillary burden and inflammatory signalling decline, the serum proteome can therefore change through reduced inflammatory consumption, altered liver synthesis, normalization of vascular permeability, tissue repair and recovery of lipoprotein and endocrine networks. The longitudinal signal is not simply an inflammatory score: its coordinated carrier and lipid components suggest that molecular recovery includes rebuilding the extracellular transport systems that connect liver, adipose tissue, immune cells and damaged lung.

BMI was the available anthropometric proxy in this reanalysis, not a direct measurement of nutritional state. The absence of false-discovery-rate-significant baseline protein differences or BMI-by-time interactions therefore does not imply that nutrition is biologically unimportant in TB. Two individuals with the same BMI may differ substantially in skeletal muscle, visceral fat, dietary intake, micronutrient stores, inflammation, oedema and hepatic function. Active TB can also alter circulating carrier proteins independently of intake, so molecular and anthropometric phenotypes need not align closely. Direct body-composition and micronutrient measurements will be required to resolve this heterogeneity prospectively.

The serum nutritional-metabolic recovery axis integrates lipid-transport, nutritional-carrier and endocrine-immunometabolic proteins into a compact summary phenotype. Because coordinated treatment-associated behaviour informed selection of the constituent modules, their shared longitudinal increase was expected to be reflected in the integrated component. The principal value of the axis is therefore not to independently demonstrate metabolic restoration, but to provide a nonredundant phenotype that can be compared across BMI strata and tested prospectively in external cohorts. Its strong correlation with the module-score construction shows that the summary signal is retained after eliminating shared-protein duplication.

These findings extend the work of Powell et al. (17) who identified 676 time-responsive proteins and showed that immune activation, haemostasis, vesicle transport, extracellular matrix and metabolism continued to change through treatment and week 52. Their observation of late apolipoprotein recovery, persistent severity-associated lipoprotein and insulin-like growth-factor signatures, and almost exclusively decreasing week-52-specific proteins is compatible with the present axis, but the analytical questions differ.

The source study asked how treatment phase, regimen and smear-defined disease severity shaped the serum proteome. The present analysis used the same longitudinal resource to test BMI-related effect modification, organize nutrition-relevant proteins into modules and reconstruct a nonredundant integrated recovery phenotype. The lack of BMI interactions therefore complements rather than contradicts the original severity findings: smear burden and BMI capture different dimensions of disease and host condition.

The module trajectories also agree with independent serum and plasma studies that have linked active TB and treatment response to acute-phase proteins, complement, coagulation, extracellular-matrix remodelling and lipid transport (16, 20). Proteomic and multi-omic studies have repeatedly identified apolipoprotein depletion or remodelling during active disease and recovery-associated changes in lipid metabolism (18, 19, 29). Differences among studies in specific proteins or direction of change are plausible consequences of population, disease stage, HIV status, specimen processing, depletion strategy, analytical platform, normalization, treatment regimen and sampling time. The present dataset is especially informative because repeated samples were obtained from the same participants through week 52; cross-sectional studies cannot separate stable interindividual differences from true within-person recovery.

Retinol, metal-handling and lysosome-related signals provide additional biological context. Retinol-binding proteins and other negative acute-phase carriers may rise as systemic inflammation subsides and hepatic transport capacity recovers. Experimental and clinical evidence links vitamin A deficiency to altered TB progression, but serum carrier abundance is not equivalent to retinol concentration or tissue vitamin A activity (13). Transferrin, haptoglobin, ferritin-associated pathways and zinc- or copper-binding proteins sit at the interface between host nutrition and nutritional immunity, through which macrophages restrict or redirect metals needed by mycobacteria (4). Lysosome/autophagy-proxy and vitamin D/humoral modules are also biologically credible because *M. tuberculosis* manipulates phagolysosomal maturation and macrophage metabolism (5, 6). Serum proteins cannot measure phagosomal acidification, autophagic flux or intracellular killing directly, but their trajectories may report systemic consequences of the cellular programmes that resolve infection and repair tissue.

The week-8 pattern merits further investigation. The preferred axis increased significantly in the BMI ≥18.5 kg/m² comparison stratum at week 8 but not in the BMI <18.5 kg/m² stratum, whereas both strata showed larger increases from week 17 onward. Formal interaction tests did not detect a BMI-dependent difference, and their confidence intervals were wide, so the early asymmetry may reflect sampling variation. A larger cohort with denser early sampling is needed to determine whether specific nutritional phenotypes show an early metabolic lag even when formal BMI-by-time interaction tests do not detect different longitudinal trajectories.

Methodologically, this reanalysis illustrates how participant-level deltas, alias-aware curation and module reduction can extract additional hypotheses from a well-characterized longitudinal proteome. The unique-protein PCA further tested whether the integrated summary depended on duplicate representation of proteins shared across modules. These approaches are transferable to other longitudinal host-response datasets, although external validation is required before the axis can be interpreted beyond this cohort.

The study has constraints that define the next experiments. The cohort was small, included only participants with durable cure and offered limited precision for interaction testing. BMI was measured at baseline, whereas longitudinal weight, body composition, dietary intake and micronutrients were unavailable. BMI strata therefore cannot establish the presence or absence of specific nutritional deficiencies, and serum carrier proteins such as retinol- or vitamin-associated proteins should not be interpreted as direct measurements of the nutrients they transport. Serum also cannot localize signals to particular tissues or immune-cell subsets, and TMT proteomics provides relative rather than absolute abundance. Future validation should combine targeted measurement of the 36-protein axis with detailed anthropometry, body composition, micronutrients, inflammatory markers, pharmacokinetics, sputum bacteriology, imaging, lung function and post-TB outcomes.

Protein-level interaction inference was sensitive to repeated-measures variance specification: the post hoc mixed-model sensitivity produced false-discovery-rate-significant interaction terms that were concentrated among sparsely paired features, whereas none of the corresponding primary paired-change contrasts survived false-discovery-rate correction. We therefore anchor protein-level inference to participant-paired change and treat the mixed-model signals as model-sensitive secondary results requiring independent validation.

Taken together, the study provides a mechanistically organized summary of longitudinal host-response biology lying between single-analyte nutritional assessment and untargeted biomarker discovery. The integrated axis should be viewed as an exploratory molecular phenotype rather than a direct nutritional-status measure. Baseline BMI was not detectably associated with the axis at baseline, while the visit-adjusted common-offset estimate was compatible with a persistent lower average level in the BMI <18.5 kg/m² stratum. Prospective studies can test whether the axis adds information when evaluated alongside direct nutritional phenotyping, treatment exposure and post-TB outcomes.

## 5. Conclusions

Longitudinal treatment of pulmonary TB was accompanied by extensive serum-proteomic remodelling, including increasing scores on an exploratory axis integrating lipid-transport, nutritional-carrier and endocrine-immunometabolic proteins. The axis increased in both baseline BMI strata, while formal BMI-by-time interaction tests did not detect differential trajectories and were too imprecise to establish equivalence. Baseline BMI was not detectably associated with the axis at baseline and should be interpreted as an anthropometric proxy rather than a comprehensive measure of nutritional status. The defined 36-protein construction provides a framework for prospective validation alongside direct measures of nutrition, drug exposure, tissue repair and long-term outcomes after TB.

## Data availability

The mass-spectrometry proteomics data generated by the source study are publicly available through ProteomeXchange/PRIDE (PXD040546) and MassIVE (MSV000091392). The present reanalysis used collaborator-supplied processed abundance, clinical metadata and TMT sample-linkage files. Participant-linked processed files are not redistributed by this manuscript. Analysis code, sanitized aggregate tables, figures, reference metadata and manuscript-supporting outputs are publicly available at https://github.com/rmaghembe1/tb-undernutrition-proteomics, with the v1.0.0 archival release available through Zenodo at DOI 10.5281/zenodo.22692544.

## Ethics statement

The source biomarker study obtained written informed consent and was approved by the University of California, San Francisco institutional review board (UCSF IRB 12-10360) (17). The present work was a secondary analysis of previously collected, de-identified processed proteomics abundance data and linked clinical and TMT sample-mapping files and involved no new participant recruitment, specimen collection, intervention, or laboratory experimentation. A separate ethics review was not sought for this secondary reanalysis. Access to the participant-linked processed files was authorized by Jon M. Jacobs and the source-study team under the existing study/data-use authorization.

## Author contributions

RSM: Conceptualization, Methodology, Data curation, Formal analysis, Investigation, Validation, Visualization, Writing – original draft. VIK, RFM, MN: Writing – review & editing. PMM, KMN: Data curation, Writing – original draft, Writing – review & editing. JMJ: Methodology, Resources. PN: Methodology, Resources. WEJ: Conceptualization, Writing – review & editing. PS: Conceptualization, Methodology. SS: Resources. SGM, SKH: Conceptualization, Supervision, Resources, Project administration. All authors reviewed and approved the final version of the manuscript.

## Acknowledgements

The authors acknowledge Powell and colleagues, the MARK-TB and REMoxTB investigators, clinical teams and participants whose work generated the longitudinal serum-proteomics resource. The authors also acknowledge the data-generating and analytical teams at the Pacific Northwest National Laboratory and collaborating institutions. The authors acknowledge the use of ChatGPT (OpenAI; version/model GPT-5.6 Sol; source: OpenAI) for language editing, structural revision, code/workflow review and submission-formatting support. All AI-assisted outputs were reviewed and verified by the authors, who take responsibility for the manuscript’s accuracy and integrity.

## Supplementary material

**Supplementary Table S1.** Dataset and analysis-output index.

**Supplementary Table S2.** Sanitized first-pass module coverage.

**Supplementary Table S3.** First-pass module summaries by visit week and BMI group.

**Supplementary Table S4.** First-pass between-BMI module-delta comparisons.

**Supplementary Table S5.** First-pass within-group paired module changes.

**Supplementary Table S6.** Curated alias-aware module coverage.

**Supplementary Table S7.** Curated module summaries by visit week and BMI group.

**Supplementary Table S8.** Curated between-BMI module-delta comparisons.

**Supplementary Table S9.** Curated within-group paired module changes.

**Supplementary Table S10.** Formal curated module longitudinal BMI-by-time interaction models.

**Supplementary Table S11.** Final Frontiers Vancouver reference-number map linking manuscript citation numbers to cited sources in first-appearance order.

**Supplementary Table S12.** Submission-package and reproducibility manifest.

**Supplementary Table S13.** Recovery-axis unique-protein membership and module overlap.

**Supplementary Table S14.** Module-level and protein-level PC1 loadings.

**Supplementary Table S15.** Within-participant recovery-axis changes from baseline.

**Supplementary Table S16.** Categorical and continuous BMI-by-time interaction models for the recovery axis.

**Supplementary Table S17.** Baseline BMI validation, score-construction correlations and final recovery-axis summary.

**Supplementary Table S18.** Covariate-sensitivity models for the serum nutritional-metabolic recovery axis.

**Supplementary Table S19.** Reconstructed longitudinal serum-proteomics dataset and analytical structure.

**Supplementary Table S20.** Ranked within-stratum protein responses during tuberculosis treatment and follow-up.

**Supplementary Table S21.** Protein mixed-model fit status.

**Supplementary Table S22.** Protein categorical BMI-group-by-week mixed-model sensitivity.

**Supplementary Table S23.** Retained proteins not structurally estimable in the categorical mixed-model sensitivity.

**Supplementary Table S24.** Curated-module categorical BMI-group-by-week mixed-model sensitivity.

**Supplementary Table S25.** Inferential calibration of mixed-model and paired-change protein estimates.

